# Xeno-nucleic Acid (XNA) 2’-Fluoro-Arabino Nucleic Acid (FANA) Aptamers to the Receptor Binding Domain of SARS-CoV-2 S Protein Block ACE2 Binding

**DOI:** 10.1101/2021.07.13.452259

**Authors:** Irani Alves Ferreira-Bravo, Jeffrey J. DeStefano

## Abstract

The causative agent of COVID-19, SARS-CoV-2, gains access to cells through interactions of the receptor binding domain (RBD) on the viral S protein with angiotensin converting enzyme 2 (ACE2) on the surface of human host cells. Systematic Evolution of Ligands by Exponential Enrichment (SELEX) was used to generate aptamers (nucleic acids selected for high binding affinity to a target) to the RBD made from 2’-fluoroarabinonucleic acid (FANA). The best selected ~ 79 nucleotide aptamers bound the RBD (Arg319-Phe541) and the larger S1 domain (Val16-Arg685) of the 1272 amino acid S protein with equilibrium dissociation constants (*K*_D,app_) of ~ 10-20 nM and a binding half-life for the RBD of 53 ± 18 minutes. Aptamers inhibited the binding of the RBD to ACE2 in an ELISA assay. Inhibition, on a per weight basis, was similar to neutralizing antibodies that were specific for RBD. Aptamers demonstrated high specificity, binding with about 10-fold lower affinity to the related S1 domain from the original SARS virus, which also binds to ACE2. Overall, FANA aptamers show affinities comparable to previous DNA aptamers to RBD and S protein and directly block receptor interactions while using an alternative Xeno-nucleic acid (XNA) platform.

## Introduction

The coronavirus SARS-CoV-2 has had a devastating impact on society that will likely continue into the foreseeable future. It is the third coronavirus (SARS-CoV and MERS being the other two) to emerge as a human pathogen in the past 17 years, raising the possibility that others will arise in the future (1, 2). Thus, the development of novel therapeutics targeting SARS-CoV-2 and new approaches that can potentially be extended to emerging or future viruses are urgently needed. Infection with SARS-CoV-2 requires interaction between the viral surface protein, spike (S), and a host “receptor” protein, angiotensin converting enzyme 2 (ACE2) (3), that is expressed on type II alveolar cells (4) and ciliated cells in the human airway epithelium (HAE) (5), making these cells potentially vulnerable to infection. Antibodies that block this interaction have been successfully used to mitigate COVID-2 infections (6–8).

In this report we describe the selection of aptamers, which are short nucleic acid-based sequences that bind with high affinity to targets, to block the interaction between the S protein and the ACE2 receptor. Aptamers have many applications including replacement of antibodies in biochemical assays (*e*.*g*. ELISA), utilization as biosensors (including a recent rapid test for SARS-CoV-2 (9)), and as tools for studying virus molecular biology, and development of antiviral drugs (10–18). Aptamers have shown potent antiviral activity and low toxicity in cell culture (14, 19–29) and they are among the most potent inhibitors of protein activity in vitro (30–32).

Aptamers are typically made from natural RNA or DNA using Systematic Evolution of Ligands by Exponential Enrichment (SELEX) (33, 34). More recently, Xeno-Nucleic Acids (XNA), which are nucleotide analogs with altered sugar, base, or phosphate backbones, have been employed in place of DNA or RNA. Aptamers, and in particular XNA aptamers, offer strong promise as therapeutics and diagnostics as they have low immunogenicity and, in the case of XNA aptamers, greater resistance to degradation (35–38). Our group was the first to produce 2’-fluoro-arabino nucleic acid (FANA) XNA aptamers to proteins (39, 40). These aptamers bind with exceptionally high affinity to targets and are completely RNase resistant. In this report, we describe the generation of FANA aptamers to the receptor binding domain (RBD) of the SARS-CoV-2 S protein that can block the interactions between the S protein and ACE2 receptor. Although this work is targeted for SARS-CoV-2, the established principles could potentially be used for other current or future viruses and the discovered aptamers have potential not only as virus inhibitors, but also in diagnostics and as biosensors.

## Materials and Methods

### Materials

The 2′-deoxy-2′-fluoroarabino-nucleotides (faATP, faCTP, faGTP, faUTP) required for FANA synthesis were obtained from Metkinen Chemistry (Kuusisto, Finland). Deoxyribonucleotide triphosphates (dNTPs) were from Roche or United States Biochemical (USB). Enzymes and buffers including *Taq* polymerase, T4 polynucleotide kinase (PNK), 10X ThermoPol buffer (Mg^2+^-free) and MgSO_4_ were from New England BioLabs. Radiolabeled ATP (γ-^32^P) was from PerkinElmer^®^. G-25 spin columns were from Harvard Apparatus. Miniprep DNA preparation kits were purchased from Qiagen. Nitrocellulose filter disks (Protran BA 85, 0.45 μm pore size and 25-mm diameter) were from Whatman. Magnetic beads for selection were from Invitrogen (Dynabeads^™^ His-Tag Isolation and Pulldown). All DNA oligonucleotides were from Integrated DNA Technologies IDT. Thermostable polymerase D4K for FANA nucleic acid production was prepared as described and stored in aliquots at −80°C (36). The C-terminal His-tagged SARS-COV-2 receptor binding domain (RBD) (Arg319-Phe541) was from RayBiotech. The C-terminal His-tagged SARS-CoV S1 protein (Met1-Arg667) was from SinoBiological. The His-tagged (Val16-Arg685) and untagged (Gln14-Arg685) SARS-COV-2 S1 proteins, cPass^™^ SARS-CoV-2 Neutralization Antibody Detection kit, and monoclonal antibody (clone ID: 6D11F2) were from GenScript^®^. All other chemicals were from VWR, Fisher Scientific, or Sigma.

### Methods

#### End-labeling of oligonucleotides with T4 PNK

DNA oligonucleotides were 5′ end-labeled in a 50 μl volume containing 10–250 pmol of the oligonucleotide of interest, 1X T4 PNK reaction buffer (provided by manufacturer), 10 U of T4 PNK and 10 μl of (γ-^32^P) ATP (3000 Ci/mmol, 10 μCi/μl). The labeling reaction was done at 37°C for 30 min according to manufacturer’s protocol. PNK enzyme was heat inactivated by incubating the reaction at 75°C for 15 min. Excess radiolabeled nucleotides were then removed by centrifugation using a Sephadex G-25 column.

#### Selection of FANA aptamers with SARS-COV-2 RBD using magnetic Dynabeads^™^

The 79 nucleotide FANA random pool starting material for SELEX containing a 40 nucleotide central random region flanked at the 5′ end by 20 nucleotides of DNA (5′-AAAAGGTAGTGCTGAATTCG-3′), and at the 3′ end by 19 nucleotides of fixed FANA sequence (5′-UUCGCUAUCCAGUUGGCCU-3’), was prepared as described previously (41). About 200 pmol of 5′ ^32^P-labeled FANA starting pool was heat to 90⁰ C then snap-cooled on ice. The material was then incubated with 20 pmol of SARS-COV-2 RBD protein that had been attached to Dynabeads^™^ using the C-terminal His-tag. Incubations were in 200 µl PBS (137 mM NaCl, 2.7 mM KCl, 8 mM Na_2_HPO_4_, and 2 mM KH_2_PO_4_, pH 7.4) for 30 min with agitation at room temperature. The beads were washed 2X with 200 µl of PBS and the bound FANA material was removed by adding 200 µl of imidazole containing buffer (300 mM imidazole, 50 mM sodium phosphate pH 8.0, 300 mM NaCl, 0.01% Tween^™^-20) to the beads and heating for 5 min at 90⁰ C, then removing the beads with a magnet. Bound FANA was recovered by precipitation with ethanol in the presence of 50 µg of glycogen. Material was reverse transcribed to DNA, amplified, and converted to FANA for another round of selection as previously described (41). The SELEX was stopped after round 8 as no further binding affinity increase was detected.

#### Sequence analysis of FANA products recovered from round 8

PCR products were prepared from FANA sequences recovered from round 8. The PCR material was cloned using a TOPO TA cloning kit from Life Technologies. DNA mini-preps were prepared, and the products were sequenced by Macrogen (Rockville, Maryland). The appropriate DNA oligonucleotide templates for some of the recovered sequences were synthesized and generation of FANA material was performed as described (41).

#### Determination of apparent equilibrium dissociation constant (*K*_D,app_) using nitrocellulose filter binding assay

Standard reactions for *K*_D,app_ determinations were performed in 20 µl of PBS with 0.1 mg/ml BSA and 0.1 nM 5′ ^32^P end-labeled aptamer. Increasing amounts of SARS-COV-2 RBD or other proteins were diluted in the above buffer and were added in amounts that approximately flanked the *K*_D,app_ value (estimated from initial experiments) for the aptamer. After 10 minutes at room temperature, the reactions were applied to a 25 mm nitrocellulose disk (0.45 µm pore, Protran BA 85, Whatman^™^) pre-soaked in filter wash buffer (25 mM Tris-HCl pH=7.5, 10 mM KCl). The filter was washed under vacuum with 3 ml of wash buffer at a flow rate of ~ 0.5 ml/sec. Filters were then counted in a scintillation counter. A plot of bound aptamer vs. protein concentration was fit to the following equation for ligand binding, one-site saturation in SigmaPlot in order to determine the *K*_D,app_: y = B_max_(x)/(*K*_D_+x) where x is the concentration of protein and y is the amount of bound aptamer.

#### Competition binding assays

Ten nM 5′ ^32^P end-labeled aptamer was incubated at room temperature in PBS with various amounts of excess unlabeled competitor at 0-, 1-, 2-, 4-, 8-, or 16-fold excess over radiolabeled labeled aptamer. SARS-COV-2 RBD or S1 protein was added to a final concentration of 10 nM. The total volume was 20 µl (in PBS). Incubations were continued for 1 hour. Samples were run over a nitrocellulose filter and washed and quantified as described above.

#### Dissociation constant (*k*_off_) and half-life (*t*_1/2_) determinations

Five nM 5′ ^32^P end-labeled R8-9 aptamer was incubated for 10 min at room temperature in 90 µl PBS with 5 nM SARS-COV-2 RBD protein. Nine µl was removed and filtered over a nitrocellulose filter and processed as described above. Unlabeled R8-9 aptamer was then added to the remaining 81 µl of sample in a volume of 9 µl such that the final concentrations of unlabeled R8-9 was 125 nM (25-fold excess over labeled aptamer). Ten µl aliquots were removed and filtered at 10, 20, 40, 60, 80, 100, and 120 min. A background control was prepared by mixing 5 nM 5′ ^32^P end-labeled R8-9 aptamer and 125 nM unlabeled aptamer in 9 ul of PBS, then adding 1 ul of SARS-COV-2 RBD protein (final concentration 5 nM) and incubated for 10 min before processing. The dissociation constant was determined by fitting the data from a plot of aptamer bound to the filter vs. time, to an equation for single 2-parameter exponential decay in SigmaPlot: y = ae^-bx^, where b is the dissociation constant (*k*_off_ in this case). The *t*_1/2_ value was determined from *k*_off_ using the following equation: *t*_1/2_ = 0.69/*k*_off_.

#### Binding inhibitions analysis

The ability of aptamers to block the association of SARS-COV-2 RBD with ACE-2 was measured with the cPass^™^ SARS-CoV-2 Neutralization Antibody Detection kit (GenScript^®^). For comparison, neutralizing monoclonal antibody (GenScript^®^, clone ID: 6D11F2) was also used. The manufacturer’s suggested protocol was followed.

## Results

### Selection of FANA aptamers against the spike receptor binding domain (RBD)

Aptamers were produced by a modified SELEX approach using mutated enzymes capable of converting between DNA and FANA (36). The RBD domain of the S proteins was chosen as the target because aptamers that directly block S protein-ACE2 receptor interactions were desired, rather than those that bind S protein in other domains that may be less likely to block receptor binding. The 223 amino acid RBD (amino acid Arg319-Phe541), comprises just a small portion of the S protein (1273 amino acids) (Fig. 1). It is part of the S1 subunit (amino acids Val16-Arg685) present on the outside of the viral envelope (42). The C-terminal His-tagged RBD domain was attached to magnetic beads for the selection process (see Materials and Methods). A total of 28 sequences were recovered from a limited number of clones after 8 rounds of selection. The recovered sequences were organized by sequence similarity into clusters using MAFFT (43). Sequences (7 total) from different clusters with diverse structures based on RNAfold analysis (44) were chosen for further testing. Filter binding assays were used for measuring the apparent equilibrium dissociation constants (K_D,app_) to the RBD protein and the larger S1 portion of the SARS-CoV-2 spike protein (Val16-Arg685). All 7 recovered sequences bound to the RBD and S1 protein with ~ 25-fold of greater binding affinity than the starting material (Table 1). Aptamers FANA-R8-9, the closely related FANA-R8-22, and FANA-R8-17 bound most strongly and FANA-R8-9 was chosen for further testing. A version of S1 without the His-tag bound with approximately the same affinity to FANA-R8-9 as tagged protein indicating that the His-tag played no role in aptamer binding. Binding of FANA-R8-9 to the S1 protein from SARS-CoV (2003 virus) was also tested (Table 1). The spike proteins from these two viruses, which both use ACE2 as a receptor, are ~ 76% identical at the amino acid level and ~ 74% identical in the RBD domain (45). The several-fold lower binding to S1 from SARS-CoV demonstrates the high specificity of FANA-R8-9. Binding of RBD and S1 to DNA aptamer selected for RBD binding (CoV2-RBD-1C) was also tested. Aptamer CoV2-RBD-1C has a reported K_D_ for RBD of 5 nM which suggests modestly tighter binding to RBD than FANA-R8-9 (46). Differences between the published results and ours may reflect the different affinity measurement techniques or different protein constructs (see Discussion).

**Table 1.** Appatent equilibrium dissociation constants (K_D_,app) for tested FANA aptamers

| <sup>a</sup>Aptamer | <sup>b</sup>RBD K<sub>D,app</sub><br>(nM) | <sup>c</sup>S1 K<sub>D,app</sub><br>(nM) | S1 no his tag<br>K<sub>D,app</sub> (nM) |
| --- | --- | --- | --- |
| <sup>d</sup>FANA-ST | <sup>e</sup>1695 ± 754 |  |  |
| FANA-R8-3 | 68.1 ± 16.9 | 46.4 ± 2.8 |  |
| FANA-R8-5 | 26.4 | 16.6 |  |
| FANA-R8-7 | 62.8 ± 8.4 | 31.2 ± 11.7 |  |
| FANA-R8-9 | 23.5 ± 4.6 | 14.7 ± 3.9 | 29.6 ± 3.9 |
| FANA-R8-15 | 43.3 ± 0.2 | 27.7 ± 5.3 |  |
| FANA-R8-17 | 29.5 ± 6.5 | 19.1 ± 7.5 |  |
| FANA-R8-22 | 21.3 | 13.3 |  |
| <sup>f</sup>S1-SARS-2003 |  | 259 ± 161 |  |
| <sup>g</sup>CoV2-RBD-1C | 872 ± 359 | 105 ± 34 |  |

a- Aptamer sequences (only the ~40 nt random regions of the ~79 nt aptamers are shown (see Methods)):
FANA-ST: 5'-NNNNNNNNNNNNNNNNNNNNNNNNNNNNNNNNNNNNNNNNNNNNNNNNNNNNNNNN-3'
b- "RBD": Receptor binding domain of SARS-CoV-2 spike protein with C-terminal His-tag (see Fig. 1).
c-“S1”: S1 portion of SARS-CoV-2 spike protein with C-terminal His-tag (see Fig. 1).
d- FANA-ST: starting material for the SELEX procedure (see text for sequence).
e- $K_{D,app}$ values were determined using nitrocellulose filter binding in PBS buffer. Results are averages of 2 -3 experiment ± standard deviation (for experiments with 3 determinations only).

**Figure 1.**
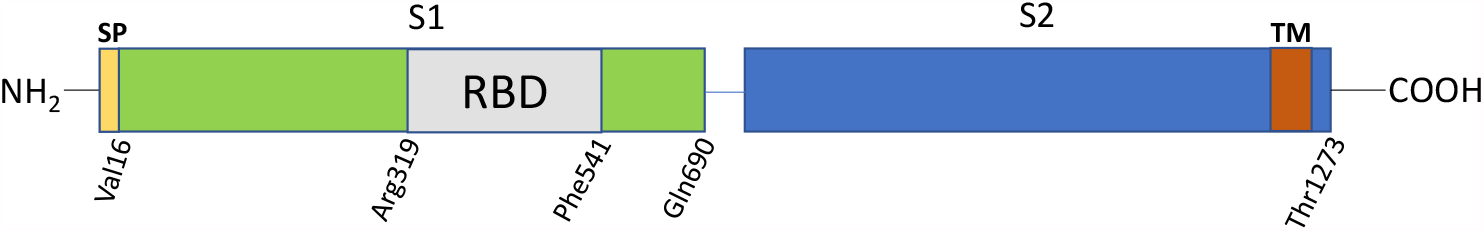
Structure of SARS-CoV-2 spike protein. The spike (S) protein has two major domains, Subunit 1 (S1) and Subunit 2 (S2). The receptor-binding domain (RBD) that binds to ACE2 is amino acid Arg319-Phe541 of the S1 domain. A 16 amino acid single peptide (SP) is present at the start of the N-terminus while the transmembrane domain (TM) is located near the C-terminus (amino acids 1213-1237).

The sequences of FANA-R8-9 and FANA-R8-17 are aligned with other recovered sequences from the same clusters (Fig. 2A) and the predicted structures of FANA-R8-9 and FANA-R8-17 are shown in Fig. 2B. The other sequences from these clusters had similar predicted structures. Aptamers from other clusters in Table 1 (FANA-R8-3, FANA-R8-7, and FANA-R8-15) that bound less tightly also had different predicted structures (data not shown). Aptamer FANA-R8-9 was used in competition binding and off-rate analysis experiments. As expected, non-labeled FANA-R8-9 was able to compete with radiolabeled aptamer for binding to RBD or S1 (Fig. 3). However, non-labeled FANA-ST was unable to displace any FANA-R8-9 aptamer, even when added at 16-fold greater amounts. This confirms that FANA-R8-9 binds to RBD much more tightly than the starting material. Off-rate analysis (39) showed that FANA-R8-9 dissociated from RBD with a half-live of 53 ± 18 minutes (Ave. 3 exp. ± S.D., Fig. 4), demonstrating stable binding.

**Figure. 2.**
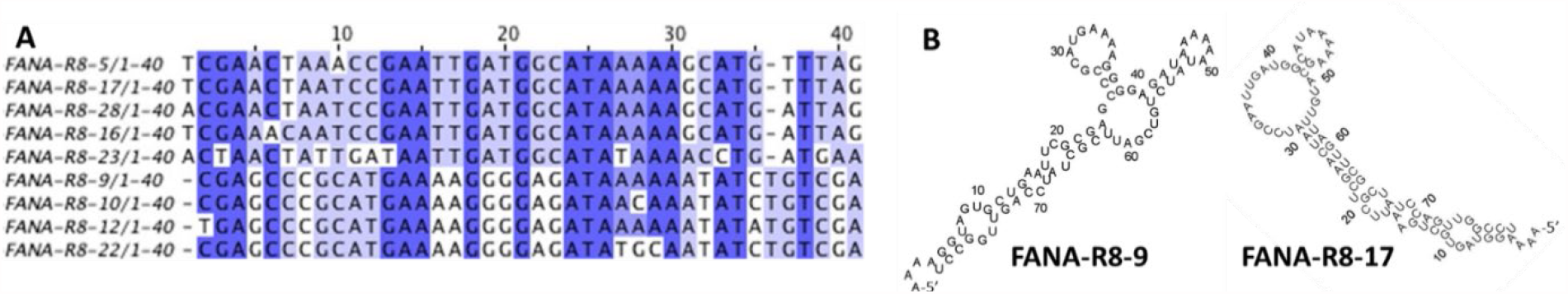
Sequence alignment using the MAFFT program for the random region (nucleotides 1-40) of recovered aptamers in the FANA-R8-17 and FANA-R8-9 lineages. MAFFT analysis of material recovered from round 8 of FANA SELEX was used to generate lineages to group related sequences. Alignments from two different lineages that contained the strongest binding aptamers are shown in (A). FANA-R8-5 to FANA-R8-23 are from the lineage containing FANA-R8-17 and FANA-R8-9 to FANA-R8-22 are from the FANA-R8-9 lineage (see panel B). The fixed primer regions: 5’-AAAAGGTAGTGCTGAATTCG-3’ at the 5’ end and 5’-UUCGCUAUCCAGUUGGCCU-3’ at the 3’ end are not shown in the alignment. (B) RNAfold program predicted structures of FANA-R8-9 and FANA-R8-17. Folded aptamers include primer regions.

**Figure 3.**
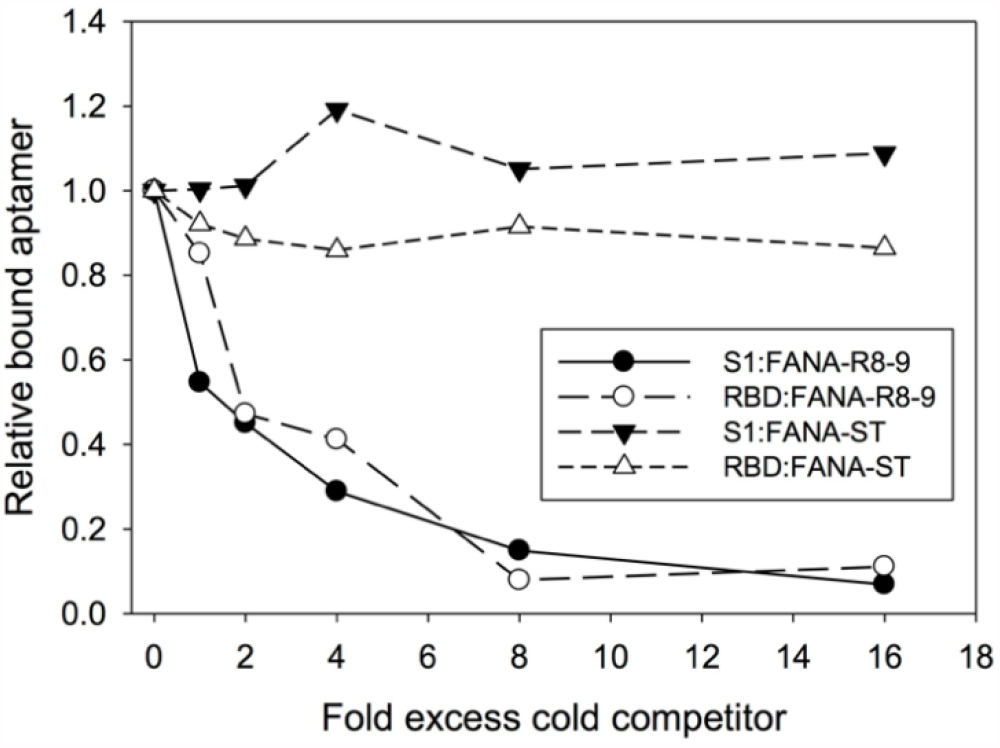
Competition binding assay with SARS-CoV-2 RBD and S1 proteins. Samples (in 20 µl PBS) contained 10 nM of 5’-^32^P end-labeled FANA-R8-9 aptamer and 10 nM of either RBD or S1 proteins. Cold competitor (FANA-R8-9 or FANA-ST (starting material for SELEX) was added at 0, 1, 2, 4, 8, and 16-fold excess over labeled FANA-R8-9. After 1-hour samples were filtered over nitrocellulose and washed with 3 ml of buffer (25 mM Tris-HCl, pH 7.5, 10 mM KCl). Filters were quantified in a scintillation counter. The background control contained no protein. The experiment was repeated with similar results. Values are relative to the value for bound radioactive material with “0” competitor.

**Figure 4.**
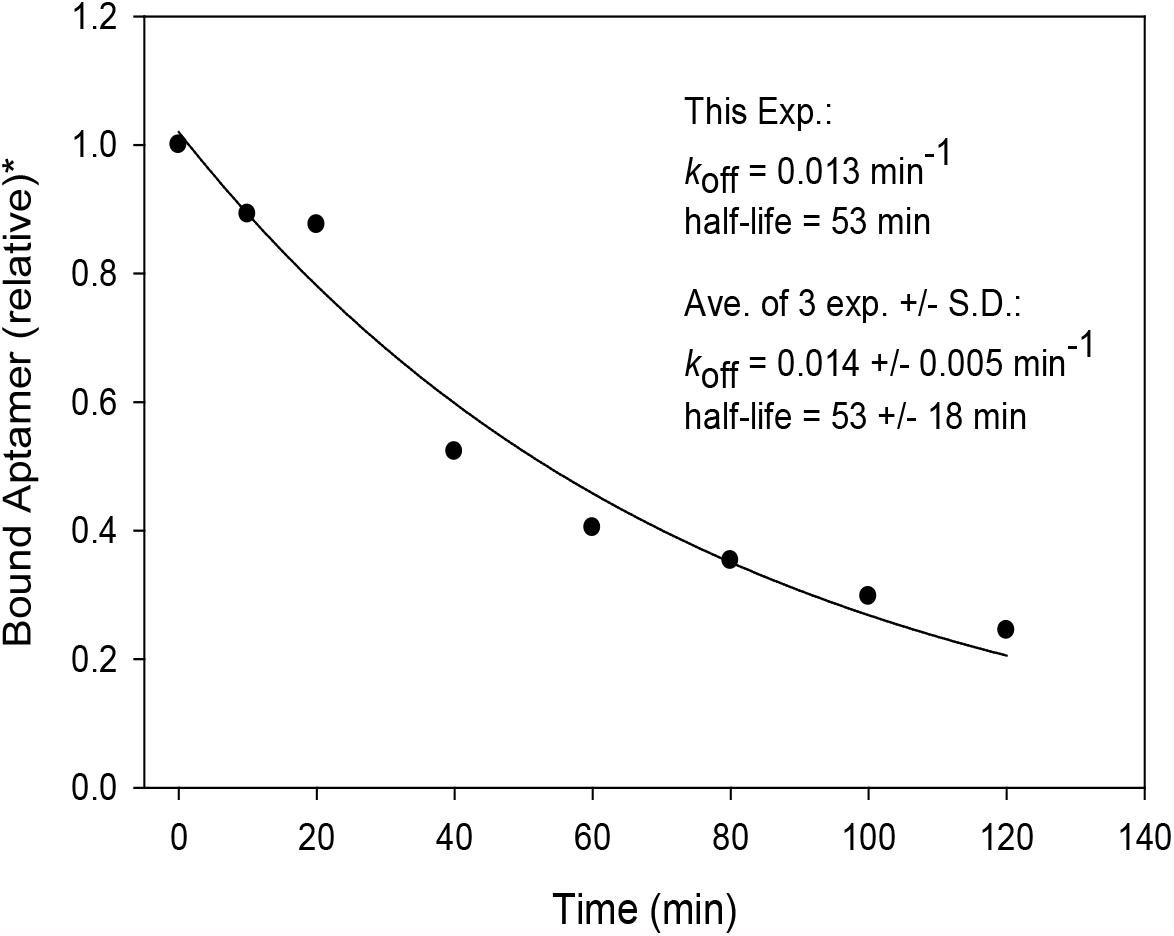
Off-rate analysis of FANA-R8-9 from SARS-CoV-2 RBD. Samples (in 90 µl PBS) contained 5 nM of 5’-^32^P end-labeled FANA-R8-9 aptamer and 5 nM RBD protein. After removal of a time “0” aliquot, 25-fold excess unlabeled FANA-R8-9 was added and aliquots were removed at 0, 10, 20, 40, 60, 80, 100, or 120 minutes and processed as described in Materials and Methods. Data was fit to a curve for single parameter exponential decay to calculate off-rate (*k*_off_) and half-life (*t*_1/2_). The experiment was repeated 3 times to yield *k*_off_ and half-life values shown. *Values are relative to the value for bound material at time 0.

### Functional assessment of lead candidate RBD-binding aptamers

To measure the ability of aptamers to block binding of the RBD to the ACE2 receptor, an ACE2 ELISA was used (Fig. 5). Aptamers were compared to an anti-SARS-CoV-2 RBD neutralizing antibody (GenScript^®^ clone ID: 6D11F2) and FANA-ST. On a weight basis, antibody 6D11F2 and FANA-R8-9 (as well as FANA-R8-22 (data not shown)) showed similar ability to block ACE2 binding (IC_50_ ~ 0.6-1.25 µg/ml) while aptamer FANA-R8-17 was ~ 3-fold weaker and FANA-ST showed no blocking of ACE2 binding (data not shown). FANA-R8-9 was also tested for stability in serum containing cell culture media (Fig. 6). Both FANA-R8-9 and CoV2-RBD-1C DNA aptamer (46) remained intact for several hours and demonstrated similar half-lives (~ 4-8 hours).

**Figure. 5.**
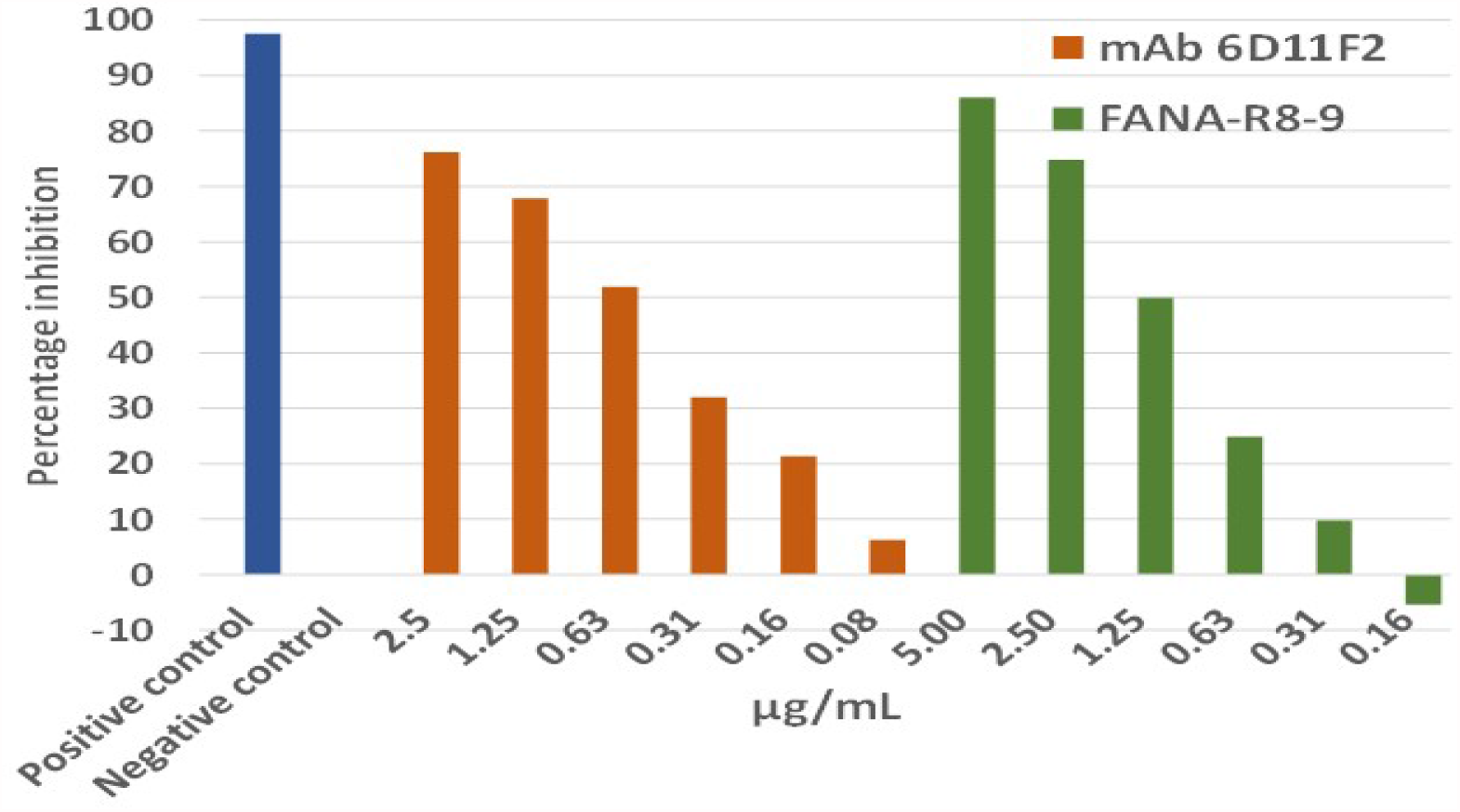
ELISA assay to test ability of antibodies and aptamers to block ACE2 binding to RBD domain. Neutralizing RBD Antibody (GenScript^®^ 6D11F2) was compared to FANA-R8-9 in an ACE2 ELISA assay (GenScript^®^). FANA-ST SELEX starting material was also tested and showed no inhibition, even at the highest concentrations (5 µg/ml) tested (data not shown). FANA-R8-17 (see Table 1) was ~ 3 times less potent than FANA-R8-9 while FANA-R8-22 was equivalent to FANA-R8-9 (data not shown). The experiment was repeated with similar results. See text for more details.

**Figure. 6.**
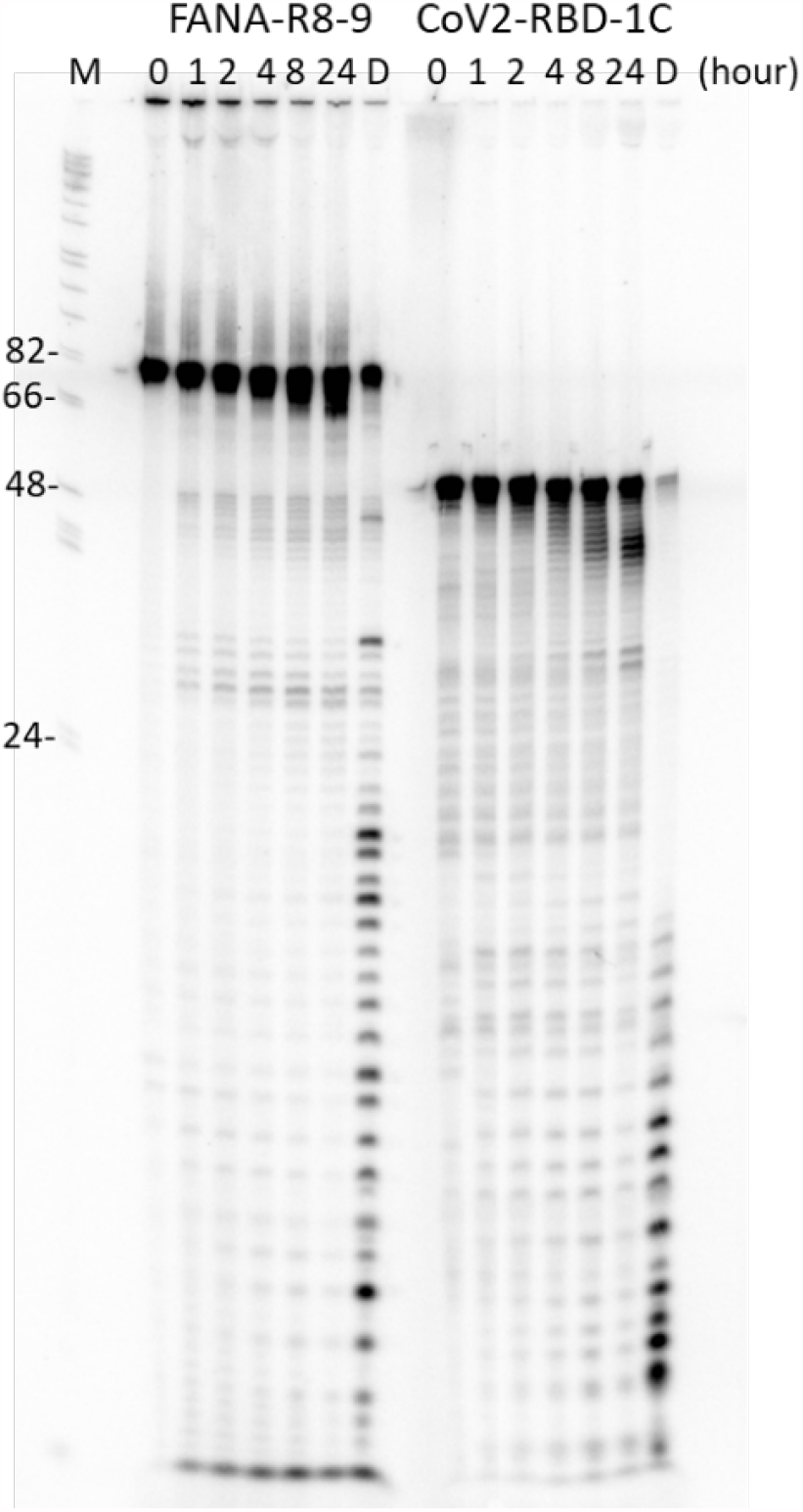
Aptamer stability assay. 100 nM of radiolabeled FANA-R8-9 (79 nts) and DNA aptamer CoV2-RBD-1C (51 nts (46)) were incubated in 200 ul of D-MEM complete + 10% FBS, and 1% penicillin/streptomycin) at 37°C. Twenty ul aliquots were removed at 0, 1, 2, 4, 8 and 24 h time points. Lane ‘D”, a 20 ul aliquot was digested with DNaseI for 30 min at 37°C as a control. The experiment was repeated with similar results.

## Discussion

This report describes the production of aptamers that can bind to and block the binding of the SARS-COV-2 RBD to ACE2. The aptamers are unique as they are made from FANA XNA as opposed to previous DNA aptamer to the SARS-CoV-2 S protein. Binding, based on K_D,app_ analysis, was comparable to previously reported DNA aptamers (46–48). The aptamers were stable for several hours in cell culture media but did break down at a rate comparable to the tested DNA aptamer (Fig. 6).

Interestingly, a previously reported DNA aptamer (Cov2-RBD-1C) that bound with a K_D_ of 5.8 ± 0.8 nM to RBD (46), did not bind strongly to the RBD in our system, although it did show binding to the S1 domain protein, albeit at a lower level than the FANA aptamers (Table 1). The RBD used for binding tests in our experiments was the same as the protein used for selection and included a C-terminal His-tag. Binding was also measured using nitrocellulose filters. The Cov2-RBD-1C aptamer was measured using RBD attached to nickel beads. It is possible that the His-tag in our measurements interfered with binding. The S1 protein used in our measurements also contained a His-tag but it is further away from the RBD domain due to the larger size of the protein. Other DNA aptamers to RBD have also been reported. Most report binding in the same low nM range as the FANA aptamers described here (46) (https://www.basepairbio.com/covid19/). This is in the same range as the reported interaction between ACE2 and the SARS-COV-2 S protein (14.7 nM), and considerably tighter than SARS S protein binding to ACE2 (325.8 nM) (49). Therefore, it would be expected that these aptamers should be good competitors for ACE2 binding. In agreement with this, FANA-R8-9 was about as effective on a per weight basis as the neutralizing RBD-specific antibody used in this analysis (Fig. 5). As there are numerous variations in the type of aptamers that can be generated with different XNAs (50, 51), perhaps those that bind even more tightly and can be obtained in the future.

The FANA-R8-9 and other aptamers (Table 1) bound with low nM affinity to RBD while previous FANA aptamers isolated in this lab to HIV RT and IN bound with low pM affinity, ~1000-fold tighter. One reason for this is RT and IN are both natural nucleic acid binding proteins and already bind tightly to specific nucleic acids. It is more of a challenge to recover strong binding aptamers to proteins that do not naturally bind nucleic acids. However, this is not always the case. Aptamers to thrombin, for example, can bind with pM affinity and modified aptamer to VEGF, which is the target for aptamer therapy for macular degeneration, also show pM binding (52–54). Several aptamers made using Slow Off-rate Modified Aptamers (SOMA) technology that includes the addition of hydrophobic groups to nucleic acids bind tightly to targets, even those that are not natural nucleic acid binding proteins (55). Still, making aptamer is a “hit or miss” proposition and there are no guarantees that aptamer which can bind more tightly than those reported here or by others can be found. Another possible advantage of XNAs other than FANA is that some types are resistant to both RNase and DNase while FANA is only resistant to the former (35–38).

Finally, we have not yet tested the FANA aptamer in virus neutralization assays. A recent report indicates that a DNA aptamer that binds to the S1 portion of the SARS-COV-2 S protein can neutralize virus entry (47). Interestingly, this aptamer did not appear to bind to the RBD and did not block virus binding to the ACE2 receptor. This suggests that even those aptamers that do not directly block binding may be able to inhibit replication.

## Acknowledgements

This work was supported by the National Institute of Allergy and Infectious Diseases (R01AI150480) to J.J.D.

